# Biodistribution Analysis of Peptide-Coated Magnetic Iron Nanoparticles: A Simple and Quantitative Method

**DOI:** 10.1101/2023.10.11.561862

**Authors:** Pavithra Natarajan, Katherine Horak, Jennifer Rowe, Joshua Lingo, John M Tomich, Sherry D Fleming

## Abstract

Biodistribution is the tracking of compounds or molecules of interest in the subject which is integral to understanding their anticipated efficacy and safety. Nanoparticles are highly desirable delivery systems which have the ability to deliver higher nucleic acid and drug payloads and they have enhanced tumor permeability due to their unique properties such as high surface area to volume ratio. Studying the biodistribution of nanoparticles is crucial to understand their effectiveness and safety in vivo, facilitate a more application driven approach for nanoparticle development which will lead to their successful translation into clinical use. In this study, we present a relatively simple method to determine the biodistribution of magnetic iron nanoparticles in mice. Branched Amphiphilic Peptide coated Magnetic Nanobeads BAPc-MNBs like their counterpart i.e., Branched Amphiphilic Peptide capsules (BAPCs) with a hollow water-filled core, are readily taken up by cells in vitro and have widespread application as a nanodelivery systems. We evaluated the BAPc-MNBs tissue distribution in wildtype mice injected intravenously (i.v.), intraperitoneally (i.p.) or orally gavaged to understand the biological interactions of the peptide nanoparticles and to further the development of branched amphiphilic peptides-based nanoparticles. BAPc-MNBs were distributed widely to various organs when injected i.v. and were eliminated from the system via the intestines in feces. The spleen was found to accumulate the highest amount of BAPc-MNBs in mice administered the NPs i.v. and i.p. while they were not absorbed into the system via oral gavage. This study not only presents a relatively simple quantification method to determine in vivo biodistribution of magnetic iron nanoparticles that can be widely applied but also demonstrates the potential of Branched Amphiphilic Peptides in the form of BAPCs or BAPc-MNBs as a delivery system.

## 1. Introduction

Nanoparticles (NPs) are highly desirable and effective as drug delivery, nucleic acid delivery and theranostic agents and find application in multiple fields such as cancer therapy ^1–3^, vaccine delivery ^4^, and gene therapy. ^5^ They have unique properties including high surface area to volume ratio which facilitates delivery of higher drug loads, higher tumor accumulation due to the enhanced permeability^6^ and retention effect. Importantly, NPs may be engineered for controlled release of drugs and targeted therapy. Although significant advances have been made in the field of nanotechnology, the translation to clinical use has only been incremental. ^7^ NPs encounter a very complex environment in vivo, which influences the tissue distribution, bioavailability, and clearance from the system. ^8, 9^ Studying the biointeractions and biodistribution will allow development of nanoparticles by an application driven approach rather than a formulation driven approach which will increase the clinical use and success rate of nanoparticles. ^10^ Besides, majority of the NPs are known to accumulate in the liver which may lead to hepatotoxicity. ^11–13^ Thus, biodistribution studies are of great importance in the growing field of nanomedicine.

The cationic Branched Amphiphilic Peptides self-assemble in an aqueous solution to form peptide bilayer delimited spherical capsules called Branched Amphiphilic Peptide Capsules (BAPCs). ^14–18^ BAPCs are promising biodegradable ^14^ nanodelivery systems that have been widely explored for delivery of dsRNA, siRNA, plasmid DNA, mRNA ^19^ and radioactive nuclides. ^15, 16, 18, 20^ Conjugating branched amphiphilic peptides to the surface of metallic nanoparticles such as gold ^21^ and iron oxide nanoparticles ^22^ expanded the applications of the peptide-based delivery system and provide opportunities to study the surface properties of the peptide bilayer delimited nanoparticles. In this article we explore the biodistribution of Branched Amphiphilic Peptide bilayer coated Magnetic NanoBeads (BAPc-MNBs) in mice. ^22^

Previous studies indicated that BAPc-MNBs do not affect viability of mammalian cells in vitro and are readily taken up via multiple endocytic routes in a time dependent manner. ^22^ Thus, BAPc-MNBs are suitable candidates for in vivo applications. The initial interactions of NPs, the overall pharmacokinetics and biodistribution depend on the method of evaluation, and the tissues evaluated. Current biodistribution evaluation methods include qualitative whole animal visualization methods such as Infrared (IR) imaging, Computed Tomography (CT), Magnetic Resonance Imaging (MRI), Positron Emission Tomography (PET) or histopathology where the tissue is harvested and analyzed for nanoparticle traces. ^7^ Histopathology conventionally uses different stains or fluorescence tracking agents and therefore is only applicable for specific types of nanoparticles and ligands. Electron microscopy obtains detailed subcellular localization of nanoparticles but has limited use in studying in vivo distribution of nanoparticles. Atomic Absorption Spectroscopy (AAS) and quantitative MRI are other methods used to determine biodistribution quantitatively. 8 Biodistribution of drug containing nanoparticles are measured by the drug load in specific tissues. However, the drug distribution may not be synonymous to the nanoparticle distribution since the drug can be released prematurely and diffuse additional to tissue sites after nanoparticle deposition. ICP-MS is a quantitative method used to measure biodistribution of nanoparticles. The nanoparticles are extracted from tissues and based on elemental analysis one can quantitatively determine the biodistribution of nanoparticles. ^23^

Previously, we demonstrated that human metastatic cervical cancer cells18 readily take up BAPCs and BAPCs could deliver an HPV-16 oncoprotein encoding DNA to successfully reduce tumor cell proliferation in mice. ^20^ Therefore, based on the evidence that BAPCs have tumor penetrating abilities and the enhanced permeability effect of NPs in general, we hypothesized that BAPc-MNBs may be taken up by tumor cells and studied their biodistribution in melanoma tumor bearing mice.

In the current study, we evaluated the BAPc-MNBs tissue distribution in wildtype mice injected intravenously (i.v.), intraperitoneally (i.p.) or orally gavaged to understand the biological interactions of the peptide nanoparticles and to further the development of branched amphiphilic peptides-based nanoparticles. We also established a new relatively facile method that does not require the use of high-end instruments for determining the biodistribution of iron oxide nanoparticles. We determined that tissue distribution varied with route of application. In addition, the NPs are released via the intestines and feces as determined in this study.

## 2. Materials and methods

### 2.1. Chemical reagents and materials

Ethanol (99% pure, Sigma, ChromSolv, Denatured ethanol), 4-(2-hydroxyethyl)-1-piperazineethanesulfonic acid (HEPES) (Acros Organics, ThermoFisher), Magnetic nanobeads (Ocean Nanotech, San Diego, CA), Trifluoroethanol (TFE) (Tokyo Chemical Industry Ltd.), 3-(2-Pyridyl)-5,6-di(2-furyl)-1,2,4-triazine-5’, 5”-disulfuric acid disodium salt (Ferene-s, Sigma Ultra), Proteinase K (Sigma-Aldrich), Matrigel Matrix® (Corning), Isoflurane (Akorn Animal Health), Dulbecco’s modified Eagle’s medium (Gibco, Sigma-Aldrich), Opti-MEM-I (Gibco, Sigma-Aldrich), L-glutamine (GlutaMAX, Sigma-Aldrich) B16F10 melanoma cell line (ATCC), pH 7.4 phosphate buffered saline (PBS) with Ca^2+^ and Mg^2+^, L-ascorbic acid (Sigma-Aldrich), acetate buffer (Glacial Acetic acid, Fisher Scientific).

### 2.2. Synthesis of Branched Amphiphilic Peptide-Magnetic Nanobeads (BAPc-MNBs)

Branched amphiphilic peptide coated – magnetic nanobeads (BAPc-MNBs) were synthesized as described in Natarajan et al. ^22^ Briefly, the magnetic nanobeads were coated with a peptide bilayer by forming one peptide layer at a time on the surface of the 50nm nanoparticles. The peptide monolayer was formed on the surface by covalently binding the cysteine residue on the C-terminus of the peptide bis-(Ac-FLIVIGSII)-KKKKK-C-CONH_2_, to the maleimide groups on the magnetic nanobead surface in 75% Ethanol: HEPES solvent to prevent assembly of the peptides into spherical capsules. After washing off the excess peptides, the second peptide layer formation on the peptide monolayer coated MNBs was promoted by adding two-fold excess of bis-(Ac-FLIVIGSII)-KKKKK-CONH_2_ and adding water. The peptides self-assemble on the surface of the nanoparticles in the presence of water due to hydrophobic interactions. After sitting for 20-30 minutes on the magnetic rack, the BAPc-MNBs were carefully collected and was concentrated on a rotavapor with a 40 °C water bath. BAPc-MNBs were re-dispersed in water alone. After overnight incubation at 4 °C the BAPc-MNBs were extruded through a sterile 0.22 μM syringe filter (Millex-GS, Millipore-Sigma). This sterilizes the BAPc-MNBs and excludes any large aggregated BAPc-MNBs. The concentration of BAPc-MNBs was adjusted appropriately after quantification by ferene-s assay as stated below, such that the same lot of BAPc-MNBs were administered as high and low dose to the mice.

### 2.3. In vivo studies

C57Bl/6 mice (Jackson Laboratory) were bred and maintained in the Division of Biology at Kansas State University. Male and female C57Bl/6 mice were kept in a 12 h light/dark cycle with constant access to rodent food and water. Four-month-old male BALB/c mice were purchased from Charles River, individually housed at NWRC and maintained on a 12 h light/dark cycle with access to standard mouse chow and other forms of enrichment such as apple slices. All procedures were approved by the Institutional Animal Care and Use Committee (IACUC) and were in compliance with the Animal Welfare Act.

### 2.4. Intravenous injection of nanoparticles in C57BL/6 mice with or without tumors

C57Bl/6 mice (n= 39) were anaesthetized using isoflurane (2-3% in oxygen) prior to the injection of 100 µL of nanoparticles dispersed in 0.15 M saline i.v. Mice were randomly assigned to one of 4 groups. The mice received either (i) low dose of BAPc-MNBs (2×10^10^), (ii) high dose BAPc-MNBs (1×10^11^), (iii) Control MNB beads (2×10^10^) without peptide coating or saline (Table 1). Each group receiving MNB consisted of 8 mice while the saline control group contained 6 mice. The deeply anesthetized animals were exsanguinated and euthanized by cervical dislocation prior to collecting the tissues (spleen, lungs, kidneys, heart, brain, intestines, thymus, tumor, blood, and urine,) either 24 h or 48 h after injection. The organs were weighed, washed in saline, snap frozen in liquid nitrogen, and stored at −80°C for further analysis. The treatments that the mice were subjected to is indicated in **Table 1**. Feces were collected every 12 h over 48 h from additional mice injected with saline (N=3), low dose (n=3) or high dose (n=3) BAPc-MNBs.

Addition C57Bl/6 mice (n=30) of the same groups were injected into melanoma tumor containing mice. B16F10 (ATCC) mouse melanoma tumor cells were cultured in Dulbecco’s modified Eagle’s medium (DMEM) supplemented with 10% OptiMEM-I®, 5% FBS, 5% NuSerum, and 2mM L-Glutamine. Two million (2 × 10^6^) B16F10 cells suspended 1:1 in Matrigel were injected sub-cutaneous on the ventral side in the thoracic region of the mice. The mice were weighed, and tumor growth monitored daily. Tumor size was measured and recorded using vernier calipers as (length × width^2^)/2. The NPs were injected between 7 to 8 days after tumor cell injection and the tumor was excised 24 h or 48 h after injection of NPs and the ex-vivo tumor volume (Length X Width X Height) was calculated.

**Table 1.** Treatment of C57BL/6 mice with magnetic nanoparticles.

| Treatment | Without tumor |  | With tumor |  | Total |
| --- | --- | --- | --- | --- | --- |
|  | 24 h | 48 h | 24 h | 48 h |  |
| Saline Control | 6 |  | 6 |  | 12 |
| $2 \times 10^{10}$ BAPc-MNBs (Low dose) | 4 | 4 | 4 | 4 | 16 |
| $1 \times 10^{11}$ BAPc-MNBs (High dose) | 4 | 4 | 4 | 4 | 16 |
| $2 \times 10^{10}$ control beads (Low dose) | 4 | 4 | 4 | 4 | 16 |
|  |  |  |  |  | 60 |

### 2.5. Oral gavage mice

Eighteen (n=18) in BALB/c mice animals were fasted overnight with access to water ad lib. Mice were lightly anesthetized with isoflurane and gavaged with high dose (2×10^11^) of BAPc-MNBs in a max volume of 200 µL. Mice were euthanized at 24 h post-gavage (n= 6), at 48 h post-gavage (n= 6) or at 48 h post-sham gavage (n=6). Tissues were removed snap frozen and shipped on dry ice to Kansas State University for analysis.

### 2.6. Intraperitoneal injected mice

For i.p. injections, wildtype Balb/c mice (n=18) were briefly restrained, appropriate landmarks identified, and the lower right quadrant sterilized with isopropyl alcohol. Using a 25–30-gauge needle, high dose (2×10^11^) BAPc-MNBs were administered to mice in a volume of ≤ 1% kg body weight (n = 12) or saline was administered to the control group (n = 6). Mice were returned to their individual cages and monitored until normal behavior was observed. Organs were harvested from each treatment group at 24 h or 48 h after administration.

### 2.7. Quantification of BAPc-MNBs by Ferene-s assay in mouse tissue

The tissues were homogenized in 0.1M tris buffered saline (TBS) containing 1% Tween-20 (Sigma-Aldrich) using a pre-programmed gentleMACS™ dissociator (Miltenyi Boitec). The tissue homogenate was collected in 1.5mL Eppendorf vials and placed on a magnetic separator (Permagen) overnight for separation and collection of magnetic nanoparticles. The magnetic beads were resuspended in DI water containing five µL of 5 mg/mL Proteinase K (Sigma-Aldrich) to digest any protein bound to the NPs. The magnetic nanobeads were aliquoted into a 96 well plate and placed on handheld magnets overnight for separation at 37 ℃. The magnetic beads were washed in DI water and quantified using the ferene-s assay as described previously in Natarajan et al. ^22^ adapted to the Ferene-s chromophoric assay described in Hedayati et al. ^24^ The feces collected were weighed in 1.5mL pre-weighed tubes and treated with 300 µL of 5N HCl followed by heating for 30 min at 95°C and 700 µL of DI H_2_O was added to each tube. The debris was spun down on a benchtop centrifuge and the supernatant was collected for Fe-content analysis. Briefly, the 3-(2-Pyridyl)-5,6-di(2-furyl)-1,2,4-triazine-5’, 5”-disulfuric acid disodium salt working solution was prepared by mixing 10 mL of 5 times working buffer, 2 g L-ascorbic acid in 11 mL 2 M Acetate Buffer, and 500 µL of 0.5 M Ferene-s in DI H_2_O (0.5 g in 2 mL water). The volume was subsequently brought up to 50 mL with DIH_2_O. Ferene-s working solution (200 μL/well) was added and allowed to incubate overnight. The iron was quantified using the standard curve generated for each 96 well plate using Microsoft Excel. The statistical analysis was performed using GraphPad Prism. The iron content was normalized using the weight of the tissue.

### 2.8. Dynamic light scattering (DLS) and Zeta potential analysis

BAPc-MNBs (50 nm) and control beads were resuspended in sterile DDI H_2_O to a final concentration of 10^9^ particles/mL. Dynamic light scattering (DLS) and zeta potential (ZP) analyses were performed for nanoparticles in 10 mm path length cuvettes (Sarstedt® Standard Cuvettes) on a Zetasizer Nano ZSP (Malvern Instruments Ltd., Westborough, MA).

### 2.9. Transmission Electron Microscopy (TEM)

Five µL of undiluted NPs samples (50 nm BAPc-MNBs) were spotted on Parafilm paper. Individual grids (Lacey F/C 200 mesh Au) were carefully placed on the surface of each NP sample for ∼ 5 min. Grids were then sequentially washed with 20 µL deionized water on the parafilm. Excess sample or water were removed by gently putting the side edge of grids in contact with Kim wipes. Grids were allowed to dry overnight at ∼ 50°C in petri dishes. For imaging, grids were mounted in specimen holders specific for TEM. Conditions for imaging were set to 25 KV on a SEM Model S-4800 (Hitachi) or adjusted occasionally according to quality of images.

## 3. Result and Discussion

**Scheme 1.**
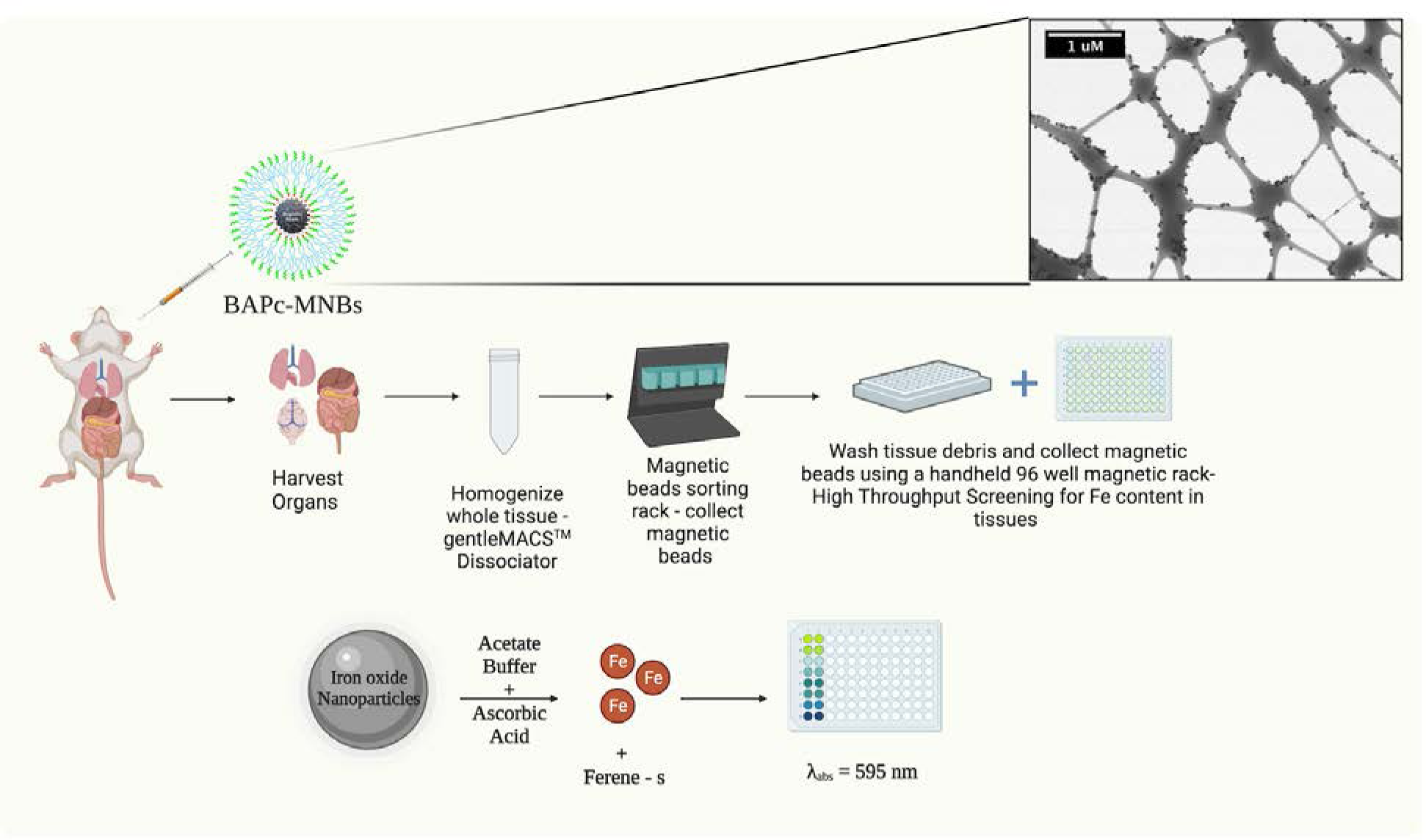
Schematic depiction of the extraction of Magnetic Nanobeads from whole organ and High throughput screening for Iron content in whole organ. (Schematic created using ©Biorender)

### 3.1. BAPc-MNBs characterization and high throughput screening of tissue for Fe content

Previously, BAPc-MNBs exhibited a hydrodynamic size of ∼200 nm by DLS analysis and approximately 50 nm by TEM analysis. ^22^ In this study, we synthesized, purified and characterized BAPc-MNBs by TEM and Dynamic Light Scattering and Zeta Potential Analysis The zeta potential and polydispersity index for BAPc-MNBs was +23 mV and <0.2, respectively, indicative of a highly homogenous solution of positively charged nanoparticles. The screening of mouse tissue for Fe was performed as described in methods and as depicted in Scheme 1.

### 3.2. Tissue distribution of in wild type C57BL/6 mice

Biodistribution of low (∼2 ×10^10^) and high dose (1×10^11^) nanoparticles administered i.v. to mice to was quantitatively assessed based on whole organ iron content at 24 h and 48 h post injection and expressed as microgram of iron detected per gram of tissue/organ. At 24 h after injection, both low and high dose BAPc-MNBs localized primarily in the spleen with higher levels at 24 h and significantly reduced levels at 48 h (**Figure 1B, D**). The average iron content detected was proportional to the dose of BAPc-MNBs injected i.e., the organs of high dosed mice had more iron per gram than the low dosed mice. The iron content ranged from ∼75 µg/g −350 µg/g in the spleen and from ∼10 µg/g −20 µg/g in the lungs and heart. Compared to saline, at 24 h the lungs, heart, and intestines contained significant NPs (**Figure 1A, C, black bars vs blue and red bars**). In addition, the intestines of mice injected with high dose BAPc-MNBs contained significantly higher iron content than saline 48 h after injection. Thus, BAPc-MNBs localized to the spleen, lungs, heart and intestines in significantly high amounts when injected i.v. in C57BL/6 mice.

**Figure 1.**
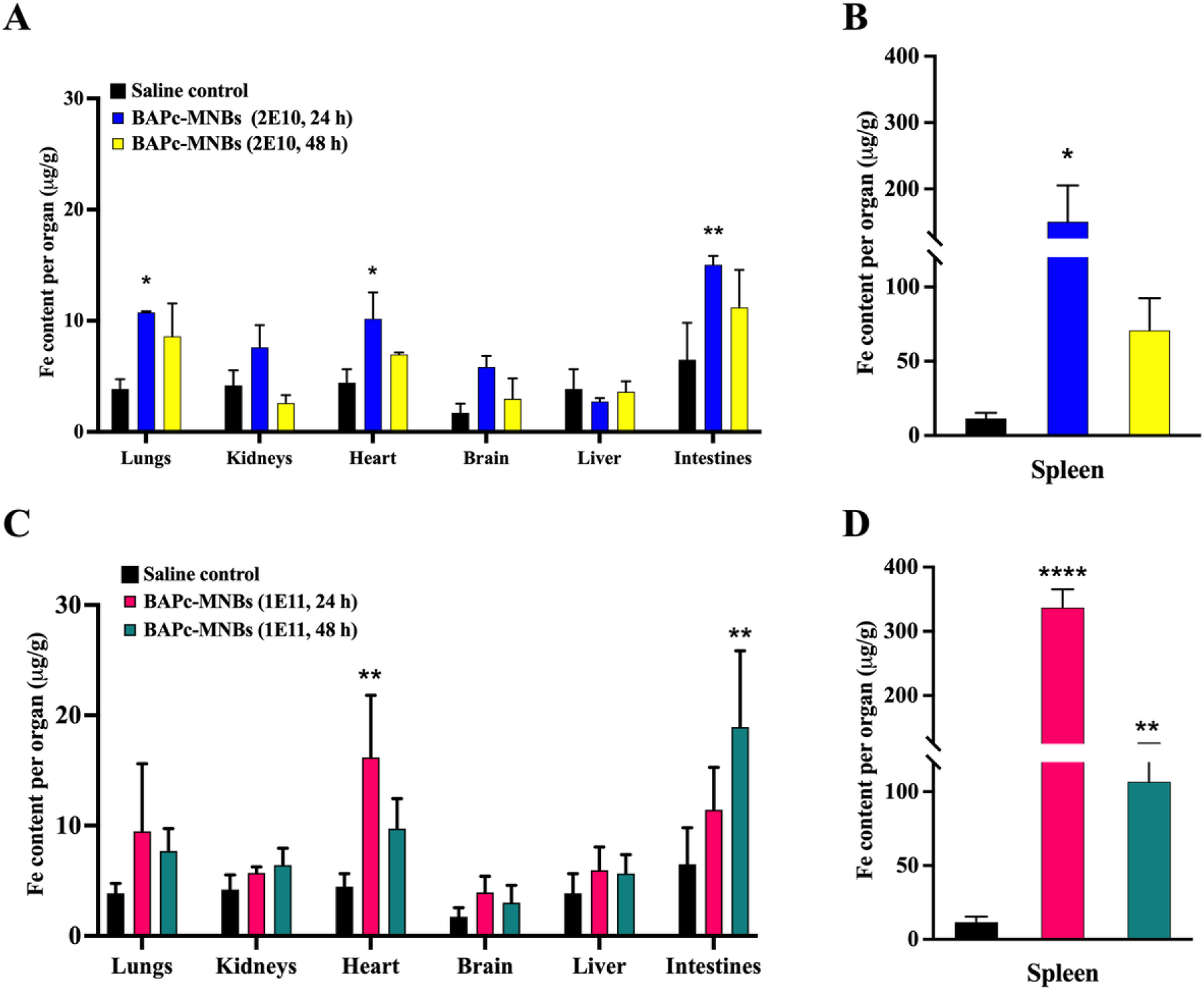
Tissue distribution of BAPc-MNBs injected i.v. in C57BL/6 mice. Fe content per gram ± SEM of whole organ harvested 24 h (blue and red bars) and 48 h (yellow and green bars) after i.v. injection with 2×10^10^ (low dose) BAPc-MNBs (A, B) and with 1×10^11^ (high dose) BAPc-MNBs (C, D). Black bars represent Fe content per gram of organs from saline control treated mice. n ≥ 3, statistical hypothesis was tested with 2-way ANOVA (A, C) and 1-way ANOVA (B, D) statistical analysis, Dunnett’s multiple comparison test. p-value: * <0.05, ** <0.01, *** <0.001, **** < 0.0001 compared to saline control.

The magnetic nanoparticles could not be detected in significant amounts in the blood (**Supplementary Figure 1**). Since the intestines showed significantly higher iron content after i.v. injection at either 24 h (low dose BAPc-MNBs) or 48 h (high dose BAPc-MNBs), it was likely that the NPs were being excreted in the feces. To test this hypothesis, feces were collected every 12 h from 3 mice per treatment group-saline control, low dose BAPc-MNBs and high dose BAPc-MNBs i.v. injected mice. The Fe content was significantly higher than saline control after 24 h of injection in mice dosed with 10^11^ (high dose) of BAPc-MNBs (**Figure 2**) and was maintained for at least 48 h after injection. This is consistent with the observed increase in iron content in the mouse intestines (**Figure 1C**) and suggests that BAPc-MNBs are eliminated via feces. By extension, it is likely that the BAPCs with the hollow water filled core may also be excreted in the feces.

**Figure 2.**
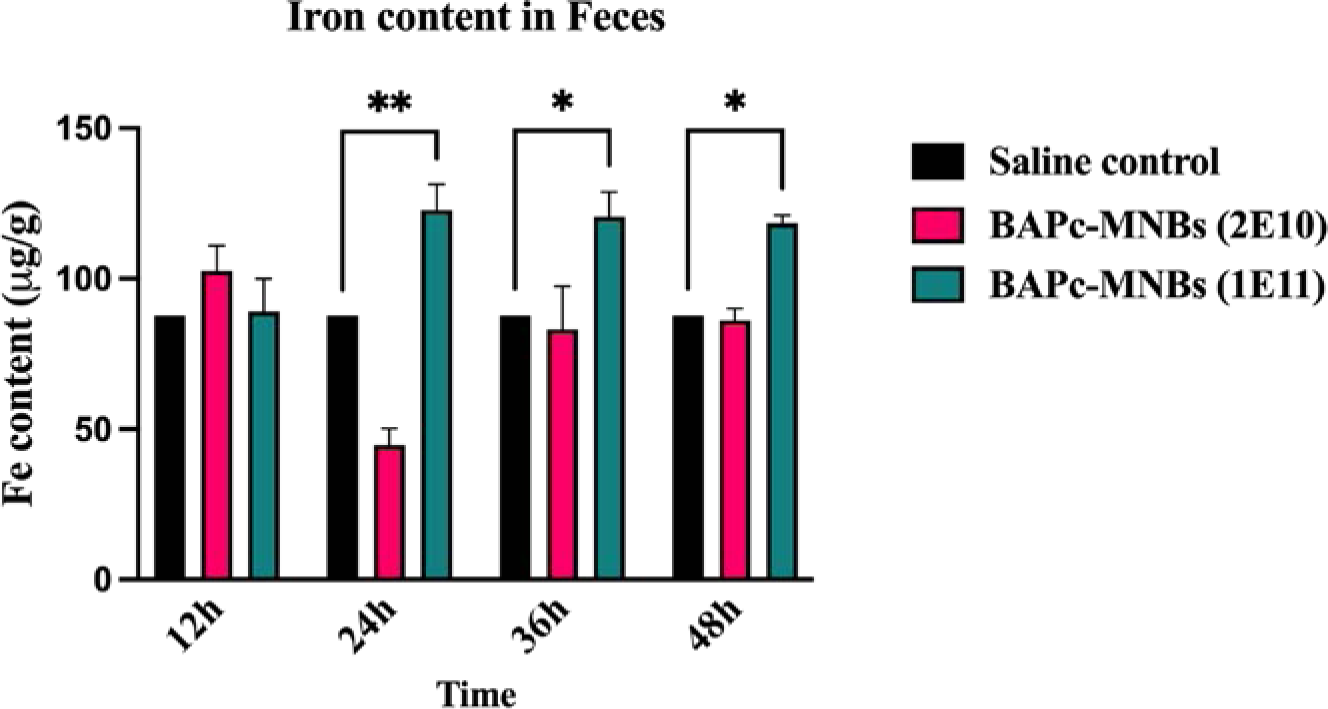
Iron content in mouse Feces. -Fe content per gram ± SEM of feces was determined in mouse feces collected at 12 h intervals. Black bars represent Fe content per gram of feces from saline control treated mice. n = 3, 2-way ANOVA statistical analysis, Tukey’s multiple comparison test for statistical hypothesis testing. p-value: * <0.05, ** <0.01, *** <0.001, **** < 0.0001

### 3.3. Tissue distribution of BAPc-MNBs in melanoma tumor bearing C57BL/6 mice

Previous studies suggested that BAPCs target tumor cells. ^18, 20^ BAPc-MNBs have the same surface chemistry as BAPCs with a hydrodynamic size of ∼200 nm. These characteristics enhance the permeability effect ^6^ increasing tumor uptake by tumors and therefore, may target tumor cells as well. To test this hypothesis, mice were injected with B16F10 melanoma cells 7-8 days prior to injecting BAPc-MNBs for determining NP tissue distribution at 24-48 h after injection. At both time points, tumor containing mice displayed a similar tissue distribution as the non-tumor bearing mice despite some changes in quantity. A significant proportion of BAPc-MNBs were localized in the spleen followed by the lungs and heart (**Figure 3 A, B, D, E**). The major difference between the tumor and non-tumor mice was the quantity of localized magnetic nanoparticles. The distribution of beads was not consistent with the dosage of NPs administered since the spleen, lungs, liver and excretory organs showed comparatively similar levels of iron content, at 24 h and 48 h after injection of low or high doses of BAPc-MNBs.

**Figure 3.**
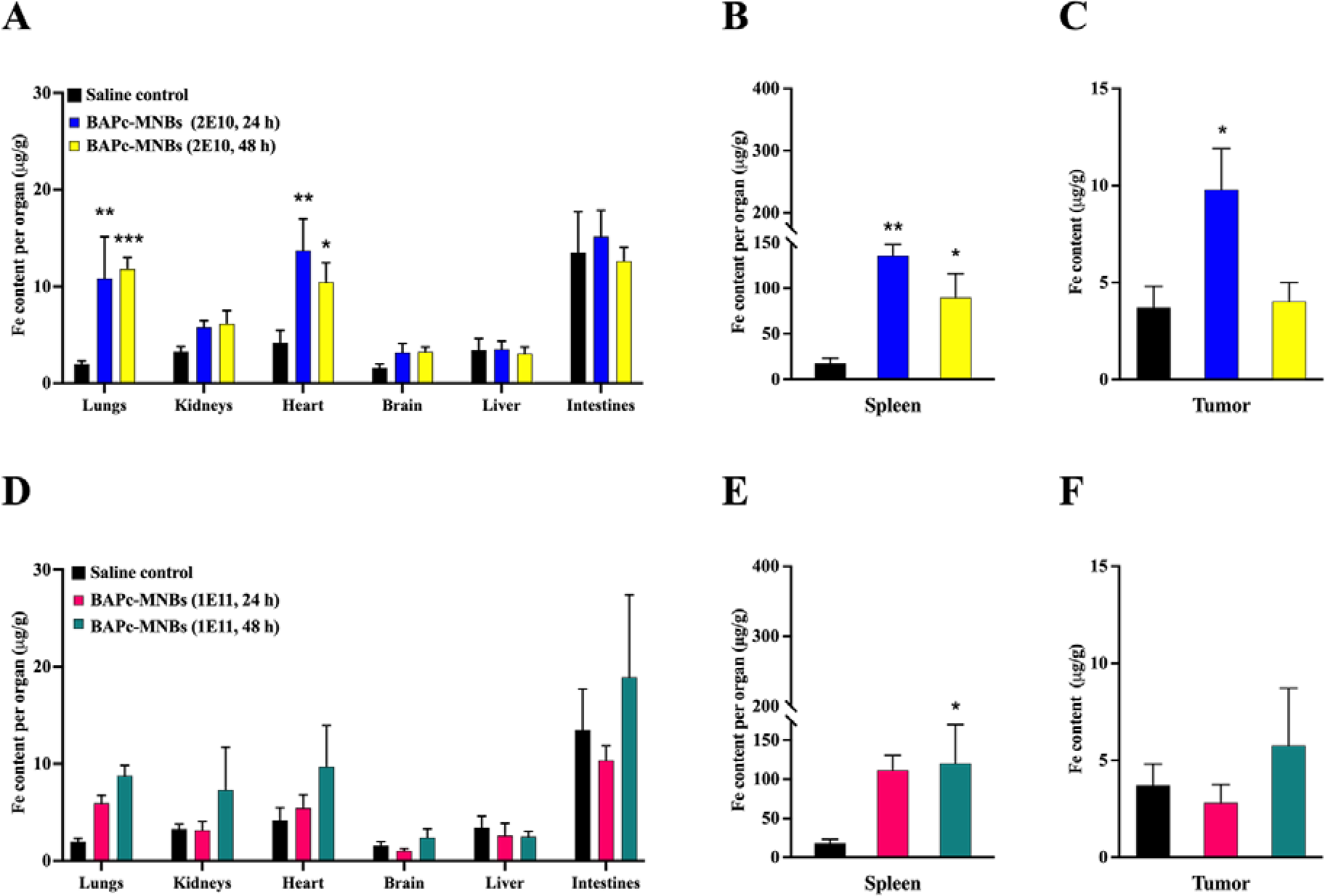
Tissue distribution of BAPc-MNBs injected i.v. in B16F10 melanoma tumor bearing mice. – Iron content ± SEM in major organs (A, D), spleen (B, E) and tumor (C, F) was determined 24 h and 48 h after injecting low dose (2×10^10^) of BAPc-MNBs and high dose (1×10^11^) of BAPc-MNBs. Black bars represent Fe content per gram of organs from saline control treated mice. n = 4, 2-way ANOVA (A, D) and 1-way ANOVA (B, C, E, and F) statistical analysis, Dunnett’s multiple comparison test for statistical hypothesis testing. p-value: * <0.05, ** <0.01, *** <0.001, **** < 0.0001

Although significantly elevated compared to saline, the spleen of mice treated with a high dose of BAPc-MNBs for 24 h showed significantly (p < 0.001) fewer NPs in the melanoma tumor bearing mice in comparison to the mice without tumors (**Figure 1, 3**). Iron content was increased in tumors harvested 24 h after treatment of mice with low dose of BAPc-MNBs only (**Figure 3C, F**).

### 3.4. Tissue distribution of BAPc-MNBs injected i.p. and administered orally in BALB-c mice

To examine the effect of route of administration, BAPc-MNBs were injected i.p. or orally gavaged into additional mice. Compared to i.v. injection, either i.p. or oral administration of BAPc-MNBs resulted significantly fewer NPs collected from all tissues. After i.p. injection, the majority of BAPc-MNBs were detected in the spleen and to some extent in the stomach and intestines (p=0.0725) after 48 h of i.p. injections (**Figure 4B, C**). Compared to the saline control, the lungs also showed increased numbers, but it was not statistically significant (**Figure 4A**). BAPc-MNBs accumulated significantly only in the spleen 48 h after administration and not detected in significant amounts in other organs tested.

**Figure 4.**
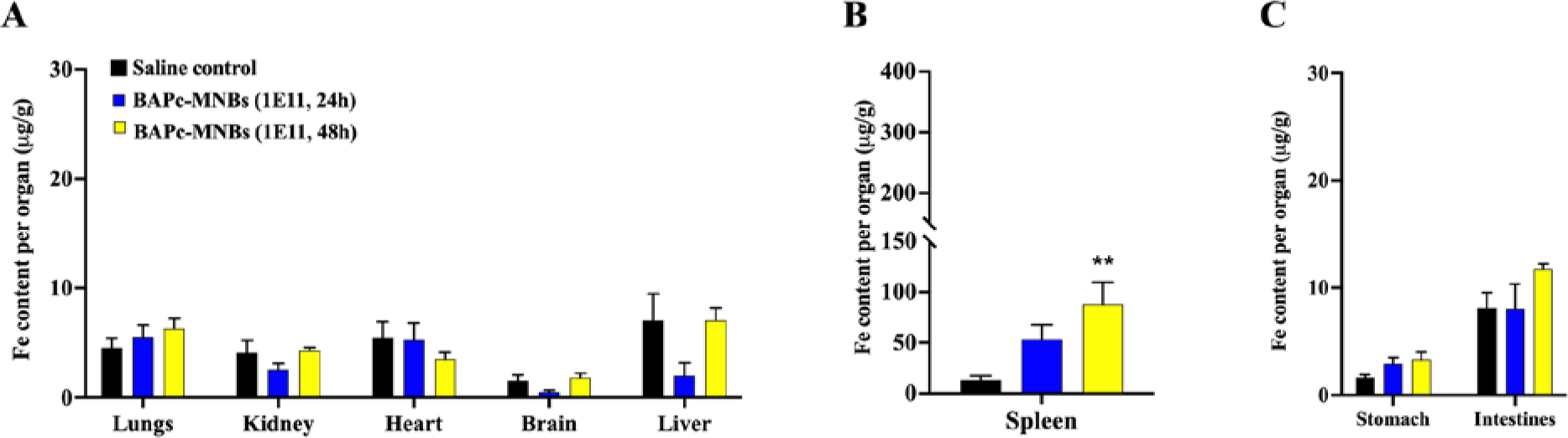
Tissue distribution of BAPc-MNBs injected IP in BALB/c mice. – Iron content ± SEM in major organs (A), the spleen (B) and the gastrointestinal system (C) was determined 24 h and 48 h after injecting high dose (2×10^11^) of BAPc-MNBs. Black bars represent Fe content per gram of organs from saline control treated mice. n = 5, 2-way ANOVA (A, C) and 1-way ANOVA (B) statistical analysis, Dunnett’s multiple comparison test for statistical hypothesis testing. p-value: * <0.05, ** <0.01, *** <0.001, **** < 0.0001

BAPc-MNBs administered orally using the gavage method were only found in significant numbers in lungs after 24 h (**Figure 5A**). No significant difference was observed between saline treated and BAPc-MNBs gavaged mice in any other major organs including the stomach and intestines. (**Figure 5A, B, C**).

**Figure 5.**
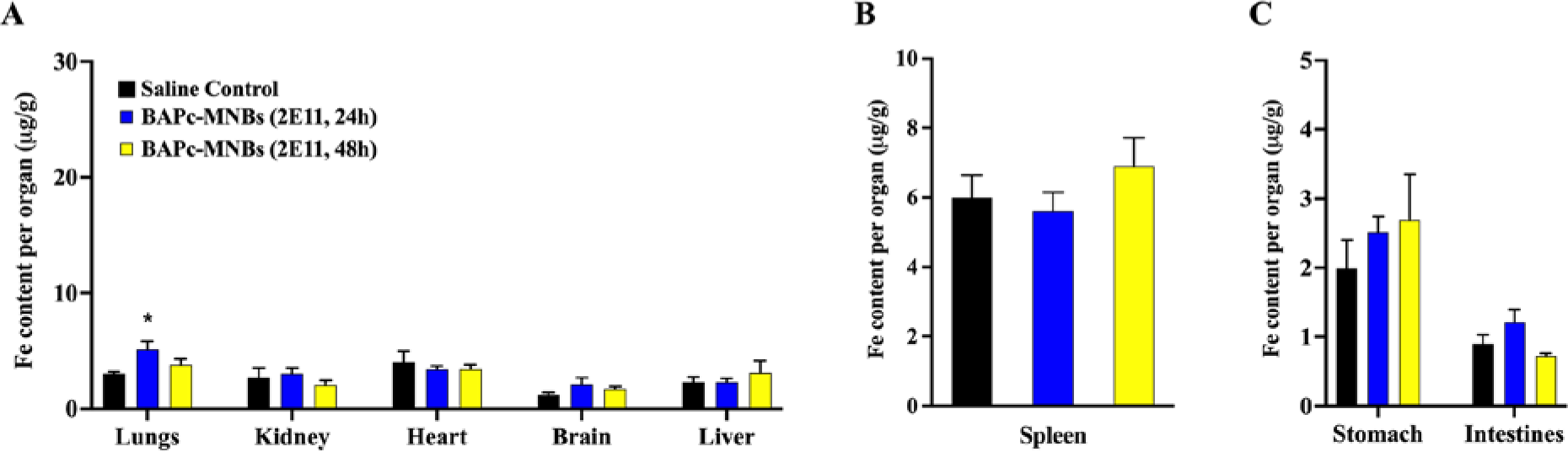
Tissue distribution of BAPc-MNBs administered orally to BALB/c mice. – Iron content ± SEM in major organs (A), spleen (B) and gastrointestinal system (C) was determined 24 h and 48 h after administering high dose (2×10^11^) of BAPc-MNBs. n = 5, 2-way ANOVA (A, C) and 1-way ANOVA (B) statistical analysis, Dunnett’s multiple comparison test for statistical hypothesis testing. p-value: * <0.05, ** <0.01, *** <0.001, **** < 0.0001.

### 3.5. Effect of Branched Amphiphilic Peptide coating on tissue distribution of magnetic nanobeads

As BAPCs appear to target tumors ^18, 20^ and changed the quantity of NPs detected in the spleen (**Figure 3B, E**) in the presence of melanoma, we hypothesized that the peptide bilayer alters the biodistribution of the MNBs. To test this hypothesis, we injected 2×10^10^ MNBs or BAPc-MNBs prior to analyzing the biodistribution. In the absence of tumors at 24 h, mice injected with control beads i.e., no peptide bilayer coating contained significantly fewer beads in the spleen (**Figure 6A**). No other organs were significantly different from control beads in the absence of tumors at 24 h or 48 h (**Figure 6A, B**). However, control beads in the intestines were significantly increased compared to saline at 24 h and 48 h after injection (**Supplementary Figure 1**)

**Figure 6.**
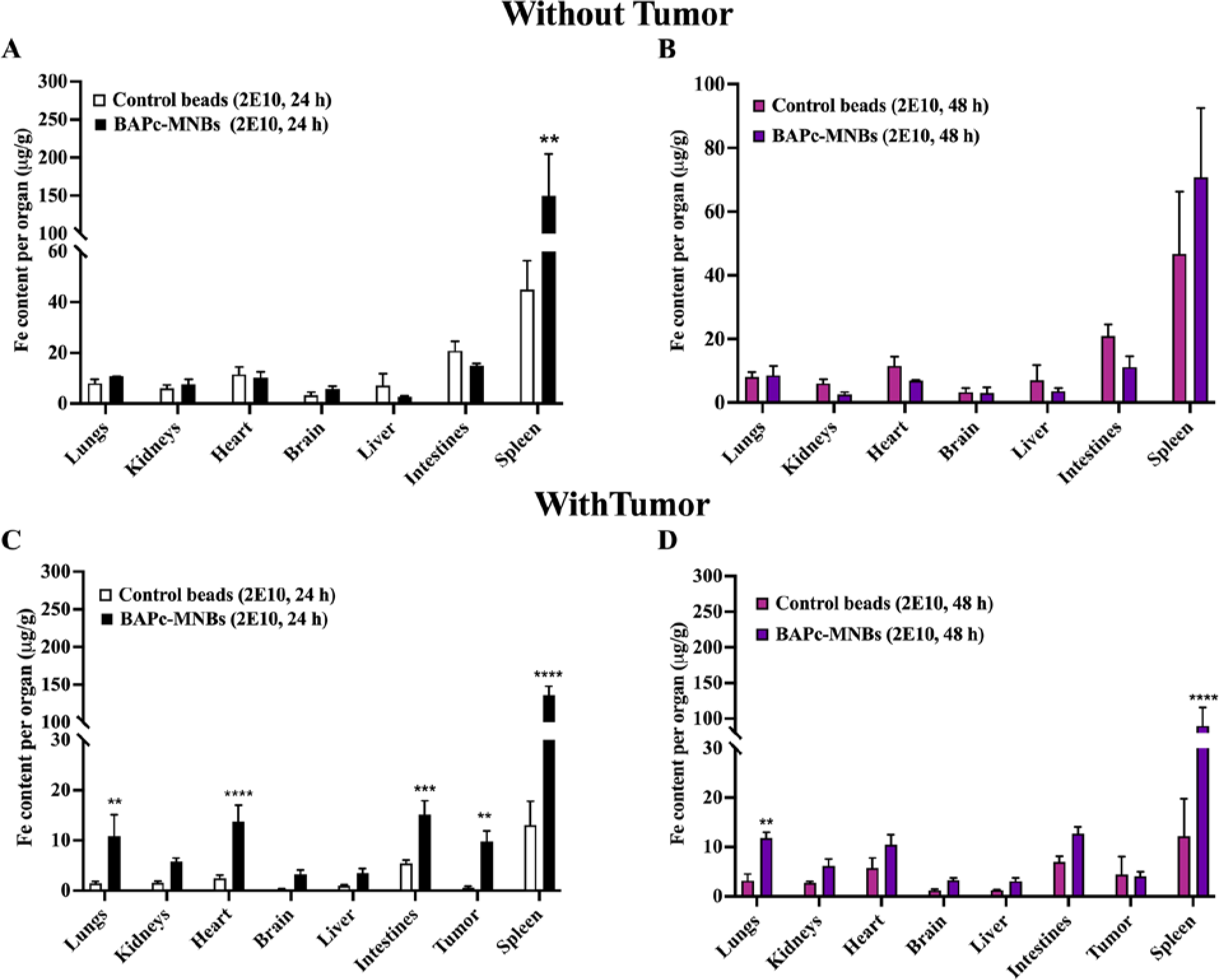
Tissue distribution of BAPc-MNBs vs control beads injected i.v. in C57BL/6 mice with or without tumors –. Fe content per gram ± SEM of whole organ harvested 24 h and 48 h after i.v. injection with 2×10^10^ (low dose) BAPc-MNBs (A, C Black; B, D purple bars) and control beads (A, C white; B, D magenta bars) in mice without (A, B) and with tumors (C, D). n ≥3, statistical hypothesis was tested with 2-way ANOVA statistical analysis, Dunnett’s multiple comparison test. p-value: * <0.05, ** <0.01, *** <0.001, **** < 0.0001 compared to saline control.

In most tissues, tumor bearing mice injected with control beads showed significantly less iron than those without tumor (**Figure 6C, D**). Similar to the findings above, the spleen contained the most control beads at both time points. The BAPc-MNBs were significantly elevated compared to control beads in the spleen of tumor bearing mice (**Figure 6C, D**) Compared to control beads, iron content was significantly increased in tumors harvested 24 h after treatment of mice with low dose of BAPc-MNBs only (**Figure 3C, D)**. The presence of melanoma tumors significantly increased the peptide bilayer containing BAPc-MNBs compared to control beads in the lungs, heart and intestines at 24 h but only the lungs remained significantly different at 48 h (**Figure 6C, D**). Thus, BAPc-MNBs localized to tumors, spleen, lungs and heart in significantly higher amounts when injected i.v. in C57BL/6 mice when compared to the control beads without the peptide coating.

## 4. Discussion

### 4.1. Development of BAPc-MNBs nanoparticles

Branched Amphiphilic Peptides Capsules are a promising nanodelivery systems which have successfully delivered nucleic acids such as eGFP encoding plasmid DNA^15^ in a variety of cell lines, anti-HPV-16 DNA vaccines ^20^ in tumor bearing mice, dsRNA^16^ and CRISPR-Cas9 in insects. The BAPc-MNBs were developed to expand the applications of the branched amphiphilic peptides. BAPc-MNBs possess the magnetic properties of the iron oxide nanoparticles and the surface properties of the branched amphiphilic peptides. In this study we determined the biodistribution of BAPc-MNBs in vivo.

A facile method was developed in this study to assess the distribution of the BAPc-MNBs in mouse organs. Briefly, the magnetic iron oxide nanoparticles were recovered from homogenized mouse organs using a rare earth magnet. The recovered nanoparticles were quantified using the spectrophotometric ferene-s assay developed by Hedayati et al. ^24^ and previously also used in our study to determine cellular uptake of magnetic nanoparticles in vitro. ^22^ We determined that the intravenously administered BAPc-MNBs were rapidly cleared from the bloodstream, accumulated mainly in spleen, lungs, heart and excreted through the intestine. In comparison to oral delivery, i.p. injections of BAPc-MNBs were slightly more successful in distributing the nanoparticles. BAPc-MNBs showed variations in tissue distribution in mice with and without subcutaneously injected melanoma tumors.

### 4.2. A simple and accurate method of determining biodistribution-ferene-s quantification

The ferene-s quantification method showed similarity to the biodistribution profiles determined by other methods such as imaging/staining ^9, 25^ and ICP-MS ^26, 27^ after i.v. injection of iron oxide nanoparticles. Sharma et al. ^28^ also showed that iron oxide nanoparticles with different surface modifications nanoparticles (CM, Dextran, PEG-PEI coated) cleared within 24 h from the bloodstream and highest concentration was observed in the spleen and liver while the lungs showed only positively charged nanoparticles (PEG-PEI coated) similar to BAPc-MNBs. The positively charged BAPc-MNBs were also observed in the lungs and spleen but the liver did not show significant number of NPs during the quantitative analysis using the ferene-s assay. It is unclear at this time if the difference is due to the surface coating or the assay.

Kunte et al. ^19^ found a similar biodistribution in mice injected i.v. with hollow water filled core BAPCs tagged with IRDye-800CW using fluorescence reflectance imaging of individual organs after euthanization or multispectral optoacoustic tomography of whole animals. BAPCs accumulated significantly in liver, lungs and spleen up to 24 h after injection. Therefore, the in vivo biodistribution profile of BAPCs and BAPc-MNBs is similar supporting the fact that NPs surface properties significantly affect their behavior in vivo. The imaging techniques used to determine BAPCs distribution support the quantitative method used in this study for BAPc-MNBs. However, the ferene-s quantitative method used in this study detected BAPc-MNBs in mice up to 48 h after injection and detected low quantities of BAPc-MNBs in lungs, heart, intestines, and feces while no detectable signal was observed using the MSOT and IR-dye based imaging techniques. Therefore, the method presented here facilitates detection of the nanoparticles in vivo quantitatively without the limitations of imaging techniques that rely on a dye or high-end imaging instrumentation.

Visual observation during the sorting of magnetic beads from the tissues clearly indicated presence of BAPc-MNBs either in the liver or the liver circulatory system. Thus, false negative results were obtained which is indicative of the limitation of this quantitative spectrophotometric assay used for analysis. No one technique is perfect and using complementary methods will help build confidence in the results obtained.

### 4.3. BAPc-MNBs biodistribution

Upon i.v. and i.p administration, BAPc-MNBs primarily localized to the spleen. Additionally significant amounts were found in the lungs, heart and intestines of i.v. injected mice. Positively charged nanoparticles upon parenteral administration are often sequestered by macrophages in the lungs, liver, and spleen.^26^ This may explain the increased number of BAPc-MNBs in the spleen. Spleen is a highly vascular organ that acts as a blood filter and receives about 4.8% of the total cardiac output. ^29, 30^ The blood carrying foreign molecules enters through the splenic artery and distributed further through a highly organized vascular system to the white pulp and the marginal zone. Once through the marginal zone about 90% of the total splenic blood flow passes through the adjacent venous sinuses continuous with the marginal zone while some enter the meshwork of the red pulp. ^29^ The marginal zone macrophages can capture BAPc-MNBs or some can be retained by the macrophages in the red pulp for slow destruction. BAPc-MNBs persist in the spleen after 24 h of injection but ∼50% reduction is observed indicating clearance and/or escape of BAPc-MNBs from the spleen by 48 h in i.v. injected mice. However, after i.p. administration an increase in iron content was observed in the spleen after 24 h from an average of 50 µg/g at 24 h to 90 µg/g at 48 h. The mouse spleen on an average weigh about 0.1 grams. The iron content in the spleen of a wild-type mouse i.v. treated with low and high dose of BAPc-MNBs for 24 h was 150 µg/g (15 µg/0.1 g) (**Figure 1B**) and 350 µg/g (35 µg/0.1g) (**Figure 1D**) which amounts to 15 µg and 35 µg net iron in the spleen, respectively. BAPc-MNBs consistently accumulated in high numbers in the spleen in comparison to all other organs at different treatment doses and times although differences were observed in the amount of NPs that accumulated over a period of time in i.v. and i.p. injected mice.

The magnetic nanoparticles could not be detected in significant amounts in the blood (**Supplementary** Figure 1). However, the significantly high iron levels observed in the intestines and feces of BAPc-MNBs treated mice (i.v.) is indicative of BAPc-MNBs being cleared from circulation within 24 h of injection and being excreted via the intestines. Due to the debris and natural Fe content in intestines, the threshold for detecting magnetic nanoparticles being excreted in comparison to saline control is much higher. Therefore, no significant increase was observed in feces of mice dosed with 2×10^10^ BAPc-MNBs. Similarly, control beads without the peptide bilayer (**Supplementary** Figure 2, **Figure 6**) were also cleared through the intestine. The enterohepatic circulation transports secreted bile from the liver to the intestines for lipid digestion and absorption of nutrients. The bile transporters are conserved between humans and mice, but the bile composition varies between the two species. ^31^ The results obtained suggests that BAPc-MNBs are transported to the intestines via the enterohepatic circulation, facilitating their clearance from the system.

Intraperitoneal injections are commonly used for treatment of localized infections and abdominal malignancies. NPs administered i.p. diffuse across the mesothelium but may not be able to diffuse across the endothelium in the connective under the mesothelial layer due to the tight junctions. However, they can enter the lymphatic system through larger openings in the peritoneum called the stomata. ^32^ BAPc-MNBs were observed in the spleen upon i.p. injection and could be distributed via the lymphatic. Not all NPs administered were detected and only a small percentage were found in the spleen and to some extent in the intestines.

NPs are not readily absorbed in the intestine of animals which is a major barrier to oral delivery. ^33^ The biocompatible molecules such as peptides and proteins are acted upon by peptidases and acidic digestive juices (pH 2) proving to be a major hurdle in oral delivery. ^34^ However, oral uptake of drugs is preferred by the end user and thus, researchers continually work to design nanoparticle-drug conjugates that are effective when administered orally. ^35^ BAPCs are unaffected by mammalian proteinases and are not denatured within mammalian cells. ^17, 18^ BAPCs conjugated with dsRNA have been successfully delivered in liquid and solid insect diets to arthropods, *Tribolium castaneum* (red flour beetle) and *Acyrthosiphon pisum* (pea aphid). ^16^ *Tribolium* shows a gradual increase from pH 5.6 to pH 7.5 in the anterior to posterior midgut, similar to mammalian intestines in pH, with the exception that their GI system is far less complex. ^36^ We therefore hypothesized that BAPc-MNBs may be able to cross the acidic stomach to be absorbed into the mammalian system through the intestinal barrier similar to their counterpart i.e., BAPCs which are highly stable under varying conditions. However, the iron content detected in all other organs was not significantly different from the control except the lung at 24 h probably due to aspiration of the NPs during gavage. This indicates that BAPc-MNBs were either present at very low undetectable levels or not present in the tissues analyzed. Thus, oral delivery using the gavage method was unsuccessful in delivering BAPc-MNBs to all the organs tested.

### 4.4. Biodistribution in presence of tumors

The presence of a subcutaneously injected melanoma tumor affected the tissue distribution of i.v. injected BAPc-MNBs and the control beads. The quantity of magnetic nanoparticles was different between the tumor and non-tumor mice and was not proportional to the dosage of NPs administered. The tumor bearing mice showed significantly lower levels of BAPc-MNBs in comparison to the non-tumor mice when dosed with high (1×10^11^) BAPc-MNBs. Kai et al. ^37^ noted that the tumor causes global immune changes which can cause faster clearance of the NPs from the system. BAPc-MNBs may have thus been cleared faster in the tumor bearing mice treated with high dose of the nanoparticles. Control beads were not present at detectable levels in the tissues analyzed, including the spleen, suggesting that they may be cleared almost instantaneously or faster than BAPc-MNBs. This is consistent with the idea that the presence of tumor accelerates the clearance of NPs. Only an average of 25% of the total administered NPs was detected in the tissues of the C57BL/6 mice, with and without tumor. The remaining NPs could be deposited in other tissues that were not analyzed, such as bones or cleared from the body. The bones have previously been demonstrated to accumulate small amounts of the actinium encapsulated BAPCs ^18^ or in lymph nodes, adipose tissue etc.

### 4.5. Effect of surface coating on biodistribution

Control beads that had no peptide bilayer coating showed some differences in the tissue distribution in comparison to BAPc-MNBs (**Supplementary Figure 2, Figure 6**). Higher amounts of control beads were observed in the liver, kidneys, and intestines compared to saline control while significantly fewer were detected in the spleen when compared to BAPc-MNBs in mice with and without tumors. The presence of tumors cleared the control beads faster from all the organs tested in comparison to BAPc-MNBs (**Figure 6**). Thus, we observed a difference in the tissue distribution of magnetic nanoparticles with the same core size but different surface composition and charge. BAPc-MNBs localized to tumors, spleen, lungs and heart in significantly higher amounts when injected i.v. in C57BL/6 mice when compared to the control beads without the peptide coating.

## 5. Conclusion

BAPc-MNBs were distributed widely to various organs when injected i.v. and were cleared from circulation within 24 h of administration. They were eliminated from the system via the intestines in feces. The spleen was found to accumulate the highest amount of BAPc-MNBs in mice administered the NPs i.v. and i.p. while they were not absorbed into the system via oral gavage. Tissue distribution is dependent on the surface chemistry due to in vivo interactions with serum proteins which leads to differences in the retention of magnetic nanoparticles in different organs.^9, 10, 26, 27^ In tumor bearing mice, control beads without the peptide coating were significantly lower than in the absence of tumors but the BAPc-MNBs appeared to target the tumor, lungs, heart, and spleen at 24 h. Thus, tissue distribution is dramatically altered early in the presence of tumors and by the peptide coating on the MNBs in the presence or absence of tumors. The method presented here to determine tissue distribution is a simple yet accurate method for quantitative analysis of iron oxide nanoparticle biodistribution administered via different routes. It can be complemented with other qualitative imaging analysis such as MRI ^38^, fluorescence live imaging ^39^ or Prussian blue staining ^9^ of tissue sections for iron content analysis. This study not only presents a relatively simple quantification method to determine in vivo biodistribution but also demonstrates the potential of Branched Amphiphilic Peptides in the form of BAPCs or BAPc-MNBs as a delivery system.

## Author Contributions

The manuscript was written through contributions of all authors. All authors have given approval to the final version of the manuscript.

## Notes

The authors declare no competing financial interests.

## ACKNOWLEDGMENT

This manuscript is contribution number 20-136-J from the Kansas Agricultural Experiment Station. We would like to thank S. Whitaker for synthesizing the peptides for our study. We would also like to thank Ms. Maria Gonzalez, the Biological Science Technician at the Electron Microscope Facility in USDA, ARS, U.S. Horticultural Research Lab, Fort Pierce, FL 34945 for the transmission electron microscopy.

## ABBREVIATIONS

BAPCs: Branched Amphiphilic Peptide Capsules, i.v., intravenously, i.p., intraperitoneally
NPs: Nanoparticles
BAPc-MNBs: Branched Amphiphilic Peptide bilayer coated Magnetic NanoBeads
IR: Infrared
CT: Computed Tomography
MRI: Magnetic Resonance Imaging
PET: Positron Emission Tomography
AAS: Atomic Absorption Spectroscopy
HEPES: 4-(2-hydroxyethyl)-1-piperazineethanesulfonic acid
TFE: Trifluoroethanol, Ferene-s,3-(2-Pyridyl)-5,6-di(2-furyl)-1,2,4-triazine-5’, 5”-disulfuric acid disodium salt
PBS: phosphate buffered saline
TBS: tris buffered saline
DLS: Dynamic light scattering
ZP: zeta potential
TEM: Transmission Electron Microscopy

---

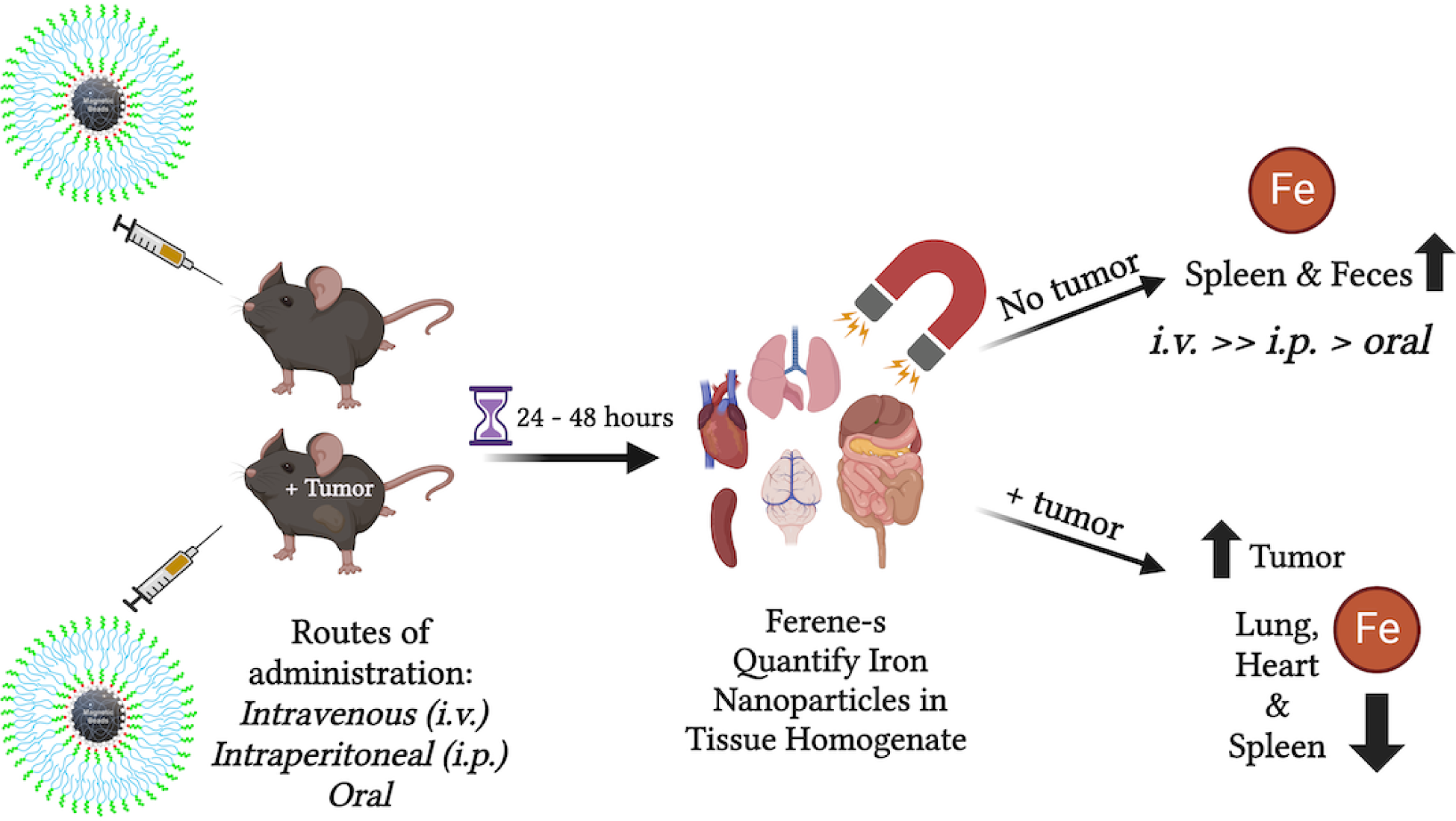
Table of Contents (TOC) Graphic.

