## Supplemental Figures 1 and 2 for "Biodistribution Analysis of Peptide-Coated Magnetic Iron Nanoparticles: A Simple and Quantitative Method"

ASSOCIATED CONTENT

Supporting Information:

Figures S1-S2

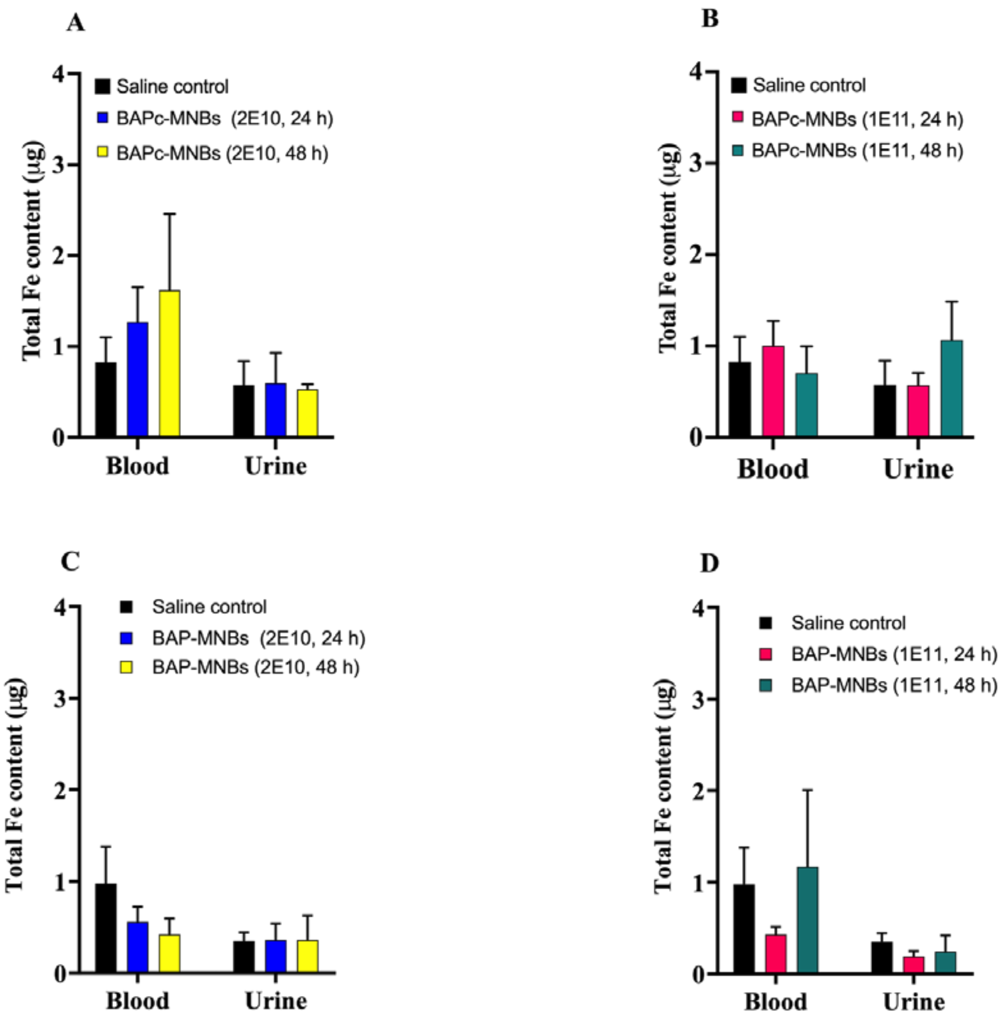


**Figure S1. Distribution of BAPc-MNBs in body fluids, injected i.v. in C57BL/6 mice** – (A), (B) without tumor and (C), (D) with tumor – Total Fe content in body fluids 24 h and 48 h after injection with low dose i.e. $2X10^{10}$ and high dose i.e. $1X10^{11}$ BAPc-MNBs. n = 3, 2-way ANOVA statistical analysis, Dunnett's multiple comparison test for statistical hypothesis testing. p-value: * <0.05, ** <0.01, *** <0.001, **** < 0.0001

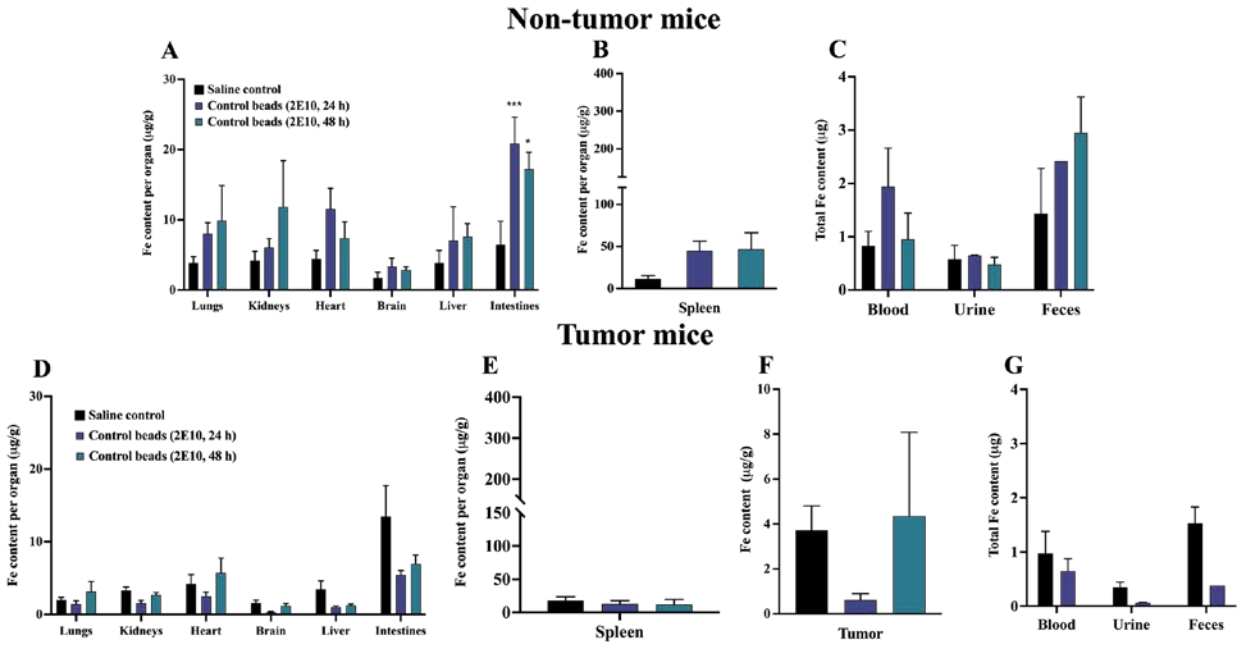


**Figure S2. Tissue distribution of control MNBs injected i.v. in C57BL/6 mice** – (A), (B), (C) without tumor and (D), (E), (F), (G) with tumor – Fe content per gram of whole organ harvested 24 h and 48 h after injection with low dose i.e. $2X10^{10}$ control MNBs i.e. with peptide bilayer coating. n = 3, 2-way ANOVA statistical analysis, Dunnett's multiple comparison test for statistical hypothesis testing. p-value: * <0.05, ** <0.01, *** <0.001, **** < 0.0001
